# Uncertainty-aware transcription factor activity and perturbation inference without additional training

**DOI:** 10.1101/2025.10.05.680506

**Authors:** Hikaru Sugimoto, Koki Tsuyuzaki, Zhaonan Zou, Shinya Oki, Tazro Ohta, Eiryo Kawakami

**Affiliations:** Department of Artificial Intelligence Medicine, Graduate School of Medicine, Chiba University, 1-8-1 Inohana, Chuo, Chiba, Chiba 260-8670, JAPAN; Predictive Medicine Special Project (PMSP), RIKEN Center for Integrative Medical Sciences (IMS), RIKEN, Yokohama, Kanagawa, JAPAN; Division of Applied Mathematical Science, RIKEN Center for Interdisciplinary Theoretical and Mathematical Sciences (iTHEMS), RIKEN, Wako, Saitama, JAPAN; Institute for Advanced Academic Research (IAAR), Chiba University, 1-33 Yayoicho, Inage,Chiba, Chiba 263-8522, JAPAN; Laboratory for Bioinformatics Research, RIKEN Center for Biosystems Dynamics Research, Wako, Saitama 351-0198, JAPAN; Institute of Resource Development and Analysis, Kumamoto University, Gene Technology Center 6F, 2-2-1 Honjo, Chuo-ku, Kumamoto-shi, Kumamoto 860-0811, JAPAN; Database Center for Life Science, Joint Support-Center for Data Science Research, Research Organization of Information and Systems, Univ. of Tokyo, Kashiwanoha-campus Station Satellite 6F. 178-4-4 Wakashiba, Kashiwa-shi, Chiba 277-0871, JAPAN

## Abstract

Understanding how transcription factors (TFs) control gene expression is essential for deciphering cellular regulatory programs. However, estimating TF activity remains challenging. Current methods rely on either broad TF–target databases, which increase false positives, or on curated sets with limited coverage. Most methods lack uncertainty quantification. Here, we introduce TFActProfiler, an integrated resource and computational framework that learns signed TF–mRNA effects by pretraining on prior TF–mRNA interaction knowledge and large RNA-seq atlases. Given an expression profile as input, TFActProfiler infers per-sample TF activity together with a confidence score that quantifies agreement between predicted and observed mRNA levels. Across TF knockdown datasets spanning multiple human cell types, TFActProfiler consistently improves TF activity inference over approaches based on unfiltered priors or narrowly curated target sets while retaining broad TF and mRNA coverage. The learned effect map further enables training-free prediction of transcriptome responses to TF perturbations. By coupling TF activity inference with training-free perturbation modeling and confidence estimates, TFActProfiler offers a tool for dissecting regulatory programs across diverse cellular states.

## Introduction

Deciphering how transcription factors (TFs) regulate gene expression has substantial translational significance beyond basic insights into cellular identity. Identifying TFs that are aberrantly activated or repressed in specific diseases can reveal mechanisms and therapeutic targets, while quantifying inter-individual variability in TF activity can yield biomarkers and enable disease subtyping (1–3). Although RNA-seq profiles thousands of transcripts across diverse contexts, expression patterns alone offer limited insight into the regulatory architecture generating these patterns. Large-scale perturbation assays, such as Perturb-seq (4), have advanced our understanding of gene expression regulation, yet comprehensive TF perturbation remains impractical in many settings. These experimental constraints have motivated the development of computational approaches that infer TF activity from transcriptomes and predict the effects of TF perturbations (3, 5–8).

Despite decades of efforts in this field (3, 5, 9–14), three challenges persist. First, achieving both wide coverage and high precision is difficult: methods maximizing coverage using co-expression or ChIP data suffer from indirect associations and non-functional binding (3, 5, 15), whereas curated regulon resources provide higher precision at the cost of limited coverage (3, 5). Second, most approaches lack uncertainty estimates despite heterogeneous and noisy evidence for TF– mRNA relationships. Third, models for predicting transcriptional responses to TF perturbations, including “foundation models” trained on large omics data (16) and regression-based models (17, 18), often require substantial task-specific training data and can sometimes fail to outperform simple baselines (6), limiting their utility in common scenarios, such as small case-control bulk RNA-seq studies without matched perturbation datasets. These limitations underscore the need for approaches that combine wide coverage with accuracy, interpretability, and applicability when matched perturbation data are unavailable.

Here, we present TFActProfiler, an integrated resource and computational framework that aims to address these challenges. This framework combines broad TF–mRNA knowledge bases with large-scale RNA-seq atlases to learn signed, quantitative TF–mRNA relationships. Across benchmarks of TF knockdowns in multiple human cell types, TFActProfiler outperforms existing methods while maintaining broad coverage of TFs and mRNAs. Our approach also provides uncertainty estimates of TF activity inference and enables predicting transcriptome responses to TF perturbations without additional training data.

## Materials and Methods

### Construction of the TFActProfiler TF–mRNA interaction resource

TFActProfiler was constructed as a resource and computational framework that learns signed TF–mRNA relationships by integrating TF–mRNA priors with large-scale RNA-seq data. This study focused on human data, taking into account the availability and quality of validation datasets. The workflow includes three steps: (I) acquisition of TF–mRNA priors, (II) acquisition of RNA-seq datasets, and (III) cluster-wise regularized regression to estimate TF–mRNA interactions.

I. Acquisition of TF–mRNA priors. Candidate TF–mRNA relationships were assembled from multiple prior resources, including ChIP-Atlas (19), the motif-predicted gene regulatory network (GRN) provided by CellOracle (17), and curated collections from CollecTRI (3). ChIP-Atlas curates public ChIP-seq, ATAC-seq, DNase-seq and bisulfite-seq datasets and provides pre-processed peak tracks. In this study, only the hg38 assembly was used. Candidate TF–mRNA pairs were extracted when the transcription start site (TSS) of the target gene was within ± 5 kb of a peak (https://chip-atlas.dbcls.jp/data/hg38/wpgsa/wpgsa_GRN.hg38.5.tsv). The human base GRN from CellOracle was obtained according to the CellOracle documentation (https://morris-lab.github.io/CellOracle.documentation/). CollecTRI-curated TF–mRNA links for humans were downloaded from decoupleR package (v2.1.1) (20). The TFActProfiler and the snapshots of the previous resources used in this study (ChIP-Atlas, the CellOracle base GRN, and CollecTRI) have been deposited in Zenodo and are available for download (10.5281/zenodo.17222986). All identifiers were harmonized to protein-coding gene symbols (GRCh38) using NCBI RefSeq (https://www.ncbi.nlm.nih.gov/datasets/genome/GCF_000001405.40/) (21). Redundant entries across sources were removed before concatenation for subsequent analyses.
II. Acquisition of RNA-seq datasets. Although these TF–mRNA priors provide potential regulatory links, the presence of binding or motif evidence does not necessarily indicate functional regulation. Additionally, it does not provide information about the mode of TF regulation (MoR; activation or inhibition). To infer MoR and quantitative effect sizes, this prior information was then combined with large-scale RNA-seq data spanning a broad range of preprocessing and biological contexts. Publicly available RNA-seq datasets were retrieved from DISCO (22, 23), Tabula Sapiens (24), the Human Cell Atlas (25–29), and ARCHS4 (30). The DISCO database provides uniformly processed, extensively curated, single-cell transcriptomic profiles. The current release contains >100 million cells across >17,000 samples, 144 tissues and 302 disease types (https://disco.bii.a-star.edu.sg/). The Human Cell Atlas is a global project that aims to describe all cell types in the human body. From the Human Cell Atlas Data Explorer (https://explore.data.humancellatlas.org/projects), Single-cell atlas of the human retina (26), The integrated Human Lung Cell Atlas (27), Human Brain Cell Atlas (28), transcriptomic cell atlas of human neural organoids (29), and endoderm-derived organoids cell atlas (25) were downloaded. The Tabula Sapiens dataset provides a cross-tissue reference for cell-type-resolved transcriptomes. This atlas includes nearly 500,000 high-quality single-cell profiles obtained from 24 tissues and organs collected from multiple donors under coordinated protocols to reduce variability arising from differences between donors. The processed data were downloaded from figshare (https://figshare.com/articles/dataset/Tabula_Sapiens_v2/27921984). ARCHS4 is a web resource that makes the majority of published RNA-seq data from human available. Gene-level expression data were downloaded from the ARCHS4 website (https://archs4.org/download).
III. Cluster-wise regularized regression to estimate TF–mRNA interactions. Each dataset was processed independently. Leiden clustering was performed at a resolution of 0.25. Clusters containing fewer than 500 cells were excluded to ensure reliable regression coefficient estimation. Within each cluster, the expression level of each target gene was modeled as a linear function of TF transcript abundances using bagging ridge regression, implemented via CellOracle (17). Ridge regression was chosen for its robustness to multicollinearity and its ability to produce stable, regularized coefficients. The default CellOracle preprocessing methods, parameters, and statistical thresholds were applied, with the exception that only less than the top 30,000 TF–mRNA pairs ranked by absolute coefficient magnitude were retained for each cluster, even if the relationship was statistically significant. Detailed preprocessing and statistical methods are provided in the CellOracle documentation (https://morris-lab.github.io/CellOracle.documentation/).

When the same pair was observed across multiple datasets or clusters, the estimated coefficients were averaged to obtain a consensus weight. Both a comprehensive global table and organ-specific tables were generated. The global table aggregated coefficients across all datasets, whereas organ-specific tables were restricted to samples annotated to the same tissue or organ system. Importantly, preprocessing heterogeneity across resources (*e*.*g*., alignment methods, quantification strategies, filtering and normalization choices already applied by the host databases) was intentionally preserved. Count matrices were used as provided by each resource, and no cross-dataset harmonization was imposed beyond the within-dataset transformations noted above. This design reflected the hypothesis that using diverse preprocessing pipelines and biological contexts at scale, analyzing each dataset under its native conditions, and averaging coefficients at the end would recover robust qualitative TF–mRNA relationships (particularly MoR and relative effect), even if exact quantitative magnitudes were not accurate and identical across datasets. Given that the strength and even MoR of TF–mRNA interactions can vary across biological contexts, the database should be interpreted as a heuristic and data-driven approximation for TF activity inference, not a definitive, universal map of transcriptional control.

### Benchmarking of TF activity inference

The benchmark quantified how well regulon-based TF activity inference identifies the perturbed TF in RNA-seq experiments, following decoupleR (v2.1.1) (20). For each perturbation with matched controls, a gene-level, signed statistic summarizing the perturbation–control contrast was supplied to enrichment procedures together with a TF–mRNA prior. The evaluation tested whether the activity score of the truly perturbed TF ranked higher than those of non-perturbed TFs in the expected direction. Benchmarking TF activity inference included four elements: (I) RNA-seq datasets in which specific TFs were perturbed and matched controls were available, (II) prior knowledge used as regulons for inference, (III) inference algorithms applied to the data and priors, and (IV) performance metrics that were used to quantify the identification of truly perturbed TFs.

I. Perturbed RNA-seq data. Perturbed RNA-seq datasets from RPE1 and K562 human cell lines (31) were used. To minimize false positives from ineffective interventions, only knockdown (KD) experiments in which the targeted TF transcript decreased relative to matched controls (fold change < 1) were retained. The pseudobulk, z-normalized matrices released with the study (31) were used directly as gene-level input. Datasets primarily targeting non-TF genes or containing too few TF perturbations (*e*.*g*., specific CRISPR screens in Adamson et al. (32)) were excluded to enable reliable estimates. Resources with potential data leakage were also excluded; for example, KnockTF (33) was not used because its literature-curated evidence overlaps with CollecTRI, creating dependence between the benchmark ground truth and one of the priors.
II. Prior knowledge. Benchmarked priors included TFActProfiler and widely used resources including ChIP-Atlas (19), base gene regulatory network provided in CellOracle (CellOracle (base)) (17), CollecTRI (3), DoRothEA (5), ChEA3-derived libraries (Literature, GTEx, ARCHS4, ENCODE, ReMap, and Enrichr) (34), five types of K562-specific regulons (S2Mb, S100Kb, S2Kb, M100Kb and M2Kb) (9), and RegNetwork (35). Given that the study (9) reported TF–mRNA relationships for K562 cells but not RPE1 cells, only the relationships for K562 cells were used. Databases such as SIGNOR (36), IntAct (37), TRRUST (38), and TFActS (39) were not evaluated separately because they are included in CollecTRI, a curated signed resource with strong performance. For each prior, signed and weighted links were used when available, and unsigned sets were treated as unit weighted.
III. Inference algorithms. Eight TF activity inference methods available in decoupleR were applied: the univariate linear model (ulm), the weighted aggregate score (waggr), and the Z-score method (zscore), univariate decision tree (udt), multivariate decision tree (mdt), gene set enrichment analysis (gsea) (40), gene set variation analysis (gsva) (41), and consensus score across methods (consensus). The algorithms are described in detail previously (20). Multivariate linear models and VIPER-like procedures (42) were not included because substantial multicollinearity in several priors produced unstable or non-convergent fits under standard settings, and prior work has shown ulm to perform better in this task (3, 20).
IV. Performance metrics. Classification performance of distinguishing the perturbed TF from non-perturbed TFs was quantified using three types of metrics, as previously described in decoupleR (v2.1.1) (20). The decoupleR documentation provides a detailed description of the methods (https://decoupler.readthedocs.io/en/latest/). First, the area under the receiver operating characteristic curve (AUROC) and the area under the precision–recall curve (AUPRC) were computed by varying a threshold on the activity scores, with the perturbed TF labeled as positive and all others as negative. A single summary was then defined as the harmonic mean (H mean) of AUROC and AUPRC. Second, an F-score was calculated from precision and recall values. Precision and recall were combined through their harmonic mean (H mean) to provide a summary F-score. Third, a quantile-rank metric (Q rank) was defined to capture the relative standing of the perturbed TF independent of absolute score scale. Scores were first ranked, quantile-normalized, and reversed by subtracting from 1, yielding values between 0 and 1, with higher values assigned to higher-magnitude scores. Perturbed and non-perturbed TFs were then compared using a one-sided Wilcoxon rank-sum test to evaluate whether perturbed TFs tended to occupy higher normalized ranks. The resulting statistic consisted of both the mean reverse-normalized rank of perturbed TFs and the associated p-value. A composite Q rank score was finally computed as the harmonic mean of the mean rank and the −log10(p-value). Finally, the individual summary metrics were aggregated into a single overall score (Score), calculated as the quantile-normalized rank of their mean.

### Reliability estimation of TF activity inference

Reliability of regulon-based TF activity inference was formalized by comparing two linear mappings between TFs and target mRNAs within each analysis unit. Let *m* and *n* denote the numbers of TFs and mRNAs, respectively, and let *c* denote the number of samples in the unit. Collect TF and gene expression into matrices *X* ∈ ℝ^*m* × *c*^ and *Y* ∈ ℝ^*n* × *c*^, whose *j*-th columns are the per-sample vectors *x*_*j*_ ∈ ℝ^*m*^ and *y*_*j*_ ∈ ℝ^*n*^.

First, a ridge-regularized regression restricted to candidate TF–mRNA links from a prior network (TFActProfiler or CollecTRI) was used to model mRNA as a linear function of TF transcripts, *Y ≈ K*·*X*, where *K* ∈ ℝ^*n* ×*m*^ is a signed coefficient matrix (rows index mRNAs, columns index TFs; coefficients outside the prior are constrained to zero). This step captures data-driven TF– gene effects under the prior’s sparsity pattern.

Second, enrichment-type activity inference was cast as a linear operator that maps mRNA back to TF activity using the same prior. For methods based on linear models or signed averaging, such as the ulm or waggr, the mapping can be written as *x*_*j*_ *≈ W*·*y*_*j*_, with *W* ∈ ℝ^*m* × *n*^ encoding signed TF–target relationships (rows = TFs, columns = mRNAs; positive entries indicate activation, negative entries repression, and missing edges are zero).

Under idealized assumptions (linearity, correct edge signs in the prior, and proportionality between TF activity and TF transcript abundance), the two mappings compose to *x*_*j*_ *≈ W*·*K*·*x*_*j*_. Deviations from this relation reflect violations of modeling assumptions or inadequacy of the prior (incorrect signs, missing edges) and therefore provide a measure of self-consistency for TF activity inference. A scale-comparable reliability score was defined as a mean-squared error (MSE) between observed TF transcripts and their reconstruction through the composed mapping. For each sample *j*, 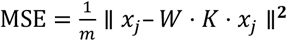. Lower values indicate higher self-consistency. The discrepancy can also be viewed as the difference between the prior knowledge of TF– mRNA relationship (*W*) and the regression-estimated TF–mRNA relationship (*K*) learned from the user-provided data.

Of note, our reliability metric should be interpreted with caution. By design, it simply quantifies the self-consistency between prior-based TF–mRNA mappings and those estimated from user-provided data. It does not diagnose which assumptions are violated when discrepancies occur (*e*.*g*., linearity, correct edge signs, or proportionality between TF activity and transcript abundance). Consequently, it serves as a heuristic, scale-comparable indicator rather than a measure of ground truth accuracy and should be considered alongside orthogonal validation as one piece of evidence.

Three case studies were analyzed. For the single cell setting, the peripheral blood mononuclear cell (PBMC) dataset distributed with decoupleR (v2.1.1) was analyzed. Megakaryocytes were excluded because only 37 cells were available, which is insufficient for stable coefficient estimation in the linear regression. For the spatial setting, a chronic active multiple sclerosis lesion dataset with Visium spot-level RNA-seq and histological annotations (43) was analyzed. Gray matter spots were excluded because only 129 spots were available. For the bulk setting, the human cancer cell line expression dataset (DepMap, Broad (2025). DepMap Public 25Q2. Dataset.) (44) was analyzed. The data were downloaded from the DepMap portal website (https://depmap.org/portal). Across all studies, TFActProfiler was compared against CollecTRI, a widely used curated resource with annotated modes of regulation. Other resources, such as ChIP-Atlas and CellOracle (base), were excluded due to their lack of annotated modes of regulation. Without signed regulatory information and in the presence of high collinearity among predictors, linear regression coefficient estimates become unstable.

### Perturbation prediction using TF–mRNA priors without additional training data

Perturbation prediction without additional training data was formulated as a linear mapping from TF transcript abundance to target mRNA under a fixed, prior-derived regulatory matrix. Let *m* and *n* denote the numbers of TFs and mRNAs in an analysis unit. Let *x* ∈ ℝ^*m*^ be the TF transcript vector and *y* ∈ ℝ^*n*^ the corresponding mRNA vector. A signed prior matrix *W* ∈ ℝ^*n*×*m*^ (*e*.*g*., from CollecTRI or TFActProfiler) encodes the expected mode of regulation (positive = activation, negative = repression; absent edges = 0). mRNA was modeled as

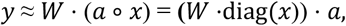

where *a* ∈ ℝ^*m*^ contains sample-specific scaling parameters that modulate each TF’s regulatory strength relative to the prior. Estimation of *a* was posed as ridge-regularized least squares after standardization. The vector *y* was standardized to have a mean of zero, and *a* was obtained as follows:

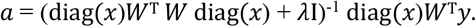

The regularization parameter *λ* was set to stabilize estimation under multicollinearity and *λ* = 0.01 was used by default. In contrast to regression frameworks that estimate an *n* × *m* parameter matrix (*e*.*g*., CellOracle-style models), the present formulation estimates only *m* parameters by fixing the interaction structure and signs through *W*. Because typically *n* ≫ *m* in transcriptomic data, the system may be constrained for stable estimation, enabling additional training data-free perturbation prediction.

After estimating the parameter vector *a*, changes in TF expression can be propagated through the model to infer the resulting changes in mRNA expression, like CellOracle, thereby predicting the transcriptional response to perturbations. For a perturbed TF *k*, the perturbed TF vector was defined as *x*^(KD)^ = *x* + Δ*x* with Δ*x* = − *γx*_*k*_*e*_*k*_, where *e*_*k*_ is the *k*-th standard basis vector and *γ* controls perturbation strength. The predicted mRNA change was then defined as follows:

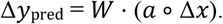

For comparisons with experimental KD RNA-seq, the observed change Δ*y*_obs_ was computed as the standardized KD-control contrast, restricted to genes present in *W*. Prediction accuracy was quantified as the Pearson correlation between Δ*y*_pred_ and Δ*y*_obs_ across genes within each perturbation experiment. In the CRISPRi setting, scientific interest lies in the direction and structure of the downstream mRNA response rather than in predicting the degree to which the TF is repressed. Therefore, the realized decrease in TF expression is included in Δ*x*. While this approach may seem to allow for data leakage, Δ*x* represents the magnitude of the administered perturbation and is considered an experimental input rather than a model target.

To evaluate the model performance, the pseudobulk, z-normalized expression matrices of perturbed RNA-seq datasets from the RPE1 and K562 human cell lines (26) were used. To minimize false positives from ineffective interventions, only KD experiments in which the targeted TF transcript decreased relative to matched controls were retained (fold change < 1). As previously described (45), the linear mapping used here should be interpreted as a first-order (local) approximation around a steady state of gene regulation. This framework also relies on the assumption that TF transcript abundance reflects TF activity, the same assumption used in CellOracle, and on the correctness and completeness of prior signs in *W*. Consistent with that interpretation, performance evaluation focused on directional agreement (*i*.*e*. correlation) rather than absolute effect size calibration. This evaluation matches our intended use in small-n bulk case–control settings, where the objective is to prioritize TF perturbations expected to move a disease transcriptome toward a healthy reference.

### Perturbation prediction with additional training data

Perturbation prediction was also evaluated in a setting where additional TF KD data were available for training. Following prior work on low-rank bilinear models (6), the expression response was decomposed into a gene embedding (*G*), a perturbation embedding (*P*), and an interaction matrix (*W*), as follows:

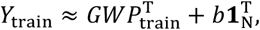

where *Y*_train_ ∈ ℝ^*g*×*N*^ is the matrix of read-out gene expression (rows = *g* genes, columns = *N* TF perturbation conditions), and *b* ∈ ℝ^*g*^ contains the row means of *Y*_train_. A *K*-dimensional gene embedding *G* ∈ ℝ^*g* × *K*^ was obtained by principal component analysis (PCA) of *Y*_train_, retaining the top *K* = 10 components as in the reference study (6). A perturbation embedding *P*_train_ ∈ ℝ^*N*×*L*^ was constructed with one row per TF perturbation (*N* TFs). Prior-driven perturbation embeddings (*P*_train_) were formed by taking the signed target vectors from TF– mRNA resources (TFActProfiler or CollecTRI), projecting them into the read-out gene space, and scaling each row by performing PCA (*L* = 50).

Additionally, embeddings derived from scFoundation (‘pos_emb.weight’) (16) were considered for comparison. In this model, *G* and *P* were derived from the embedding. Although other methods such as scGPT (46), scBERT (47), Geneformer (48), and GEARS (49) have been reported, previous benchmarks using the same RPE1 and K562 datasets found their accuracy to be consistently lower than that of the scFoundation method with this linear model (6). Accordingly, only this model was included in the present evaluation.

The interaction weight matrix *W* ∈ ℝ^*K*×*L*^ was estimated by ridge regression,

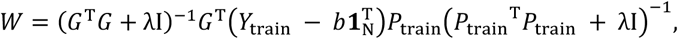

with *λ* = 0.1. After training, the expression response of a held-out perturbation with embedding vector 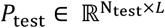 was predicted as

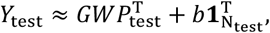

where *Y*_test_ is the predicted expression profile of the read-out genes.

For each held-out perturbation *j*, the predicted expression change 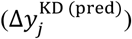 was defined as follows:

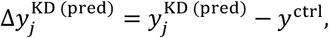

where 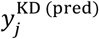 is the *j*-th column of *Y*_test_ and *y*^Ctrl^ is the matched control replicates. The observed differential-expression vector for perturbation 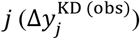 was defined as follows:

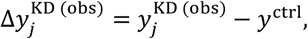

where 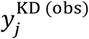 is the pseudobulk expression vector across KD replicates for TF *j*. Ten-fold cross-validation was conducted to prevent data leakage across conditions of the same TF. In each fold, *b* and *G* were recomputed using only the training split. Accuracy was quantified as the Pearson correlation between 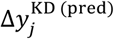 and 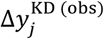, restricting to genes represented in the model (*i*.*e*., those present in *G* and *P*). Given that this model represents a local linear approximation around a steady state, emphasis was placed on directional agreement rather than absolute effect-size calibration.

### Implementation of TFActProfiler

All analyses were conducted in Python and implemented using scikit-learn (v1.7.1), NumPy (v2.2.6), Pandas (v2.3.1), Statsmodels (v0.13.2), and SciPy (v1.15.3). The TFActProfiler Python library was developed and distributed with versioned code and example notebooks (https://github.com/HikaruSugimoto/tfactprofiler). A web application for estimating TF activity from transcriptomic data (https://tfestimatetest.streamlit.app/) was developed in Python using Streamlit (https://www.streamlit.io) and was deployed on the Streamlit Cloud.

## Results

### Construction of the TFActProfiler signed TF–mRNA resource

To systematically characterize TF regulation, we integrated candidate TF–mRNA relationships from ChIP-Atlas (19), motif-based predictions (17), and curated databases (3). Since these sources often contain indirect/ambiguous links and lack quantitative information about regulatory strength, we combined them with large-scale RNA-seq atlases and linear regression models to estimate the signed influence of each TF on its downstream targets (**Figure 1A**).

**Figure 1.**
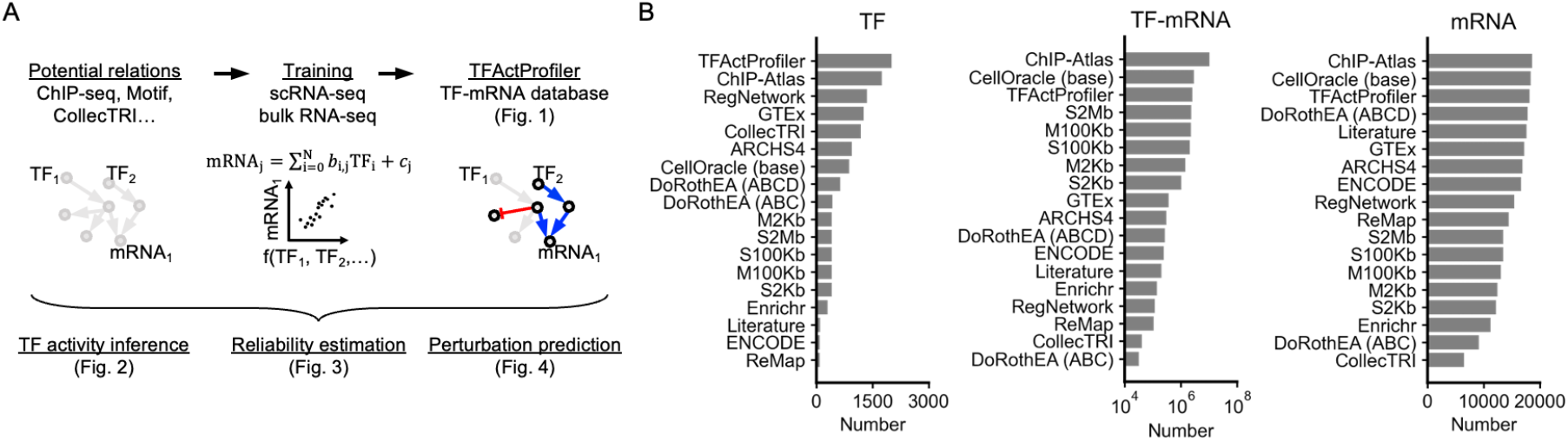
Construction of TFActProfiler, a database of transcription factor (TF)–mRNA regulatory relationships. (**A**) Overview of the TFActProfiler pipeline. Candidate TF–mRNA edges from ChIP-seq, TF binding motifs, and curated knowledge bases were integrated with large-scale bulk and single-cell RNA-seq atlases to learn signed regulatory effects (activation or repression). The resulting resource enables TF activity inference, confidence scoring, and prediction of transcriptional responses to perturbations. (**B**) Database coverage. Bar plots show the number of TFs, TF–mRNA pairs, and downstream mRNAs included in each database.

The resulting TFActProfiler database contains 2,009 TFs, 2,605,748 TF–mRNA relationships, and 18,158 target mRNAs (**Figure 1B**). Of note, according to AnimalTFDB (50), ChIP-Atlas contains 370 cofactors, CellOracle contains 31, and CollecTRI contains 198. Consequently, TFActProfiler contains 394 cofactors, and it is not limited to TFs. Compared with existing signed resources such as CollecTRI and DoRothEA, TFActProfiler includes substantially more TF– mRNA links while retaining information about the mode of regulation. Although resources like ChIP-Atlas or CellOracle’s base networks list a greater number of potential associations, some of these links do not correspond to functional regulation and lack information about the mode of regulation. In contrast, TFActProfiler integrates evidence for the mode and strength of regulation, making it one of the most comprehensive and functionally informative TF–mRNA regulatory databases to date.

### TF activity inference using TF–mRNA priors

We then benchmarked the performance of TFActProfiler in inferring TF activity using previously established strategies (20). We evaluated regulon-based TF activity inference using RNA-seq perturbation datasets in which specific TFs were knocked down (KD) (**Figure 2A**). The task for each experiment was to prioritize the perturbed TF from among all regulators using activity scores derived from a TF–mRNA prior. The priors included TFActProfiler, ChIP-Atlas (19), the base gene regulatory network provided in CellOracle (CellOracle (base)) (17), CollecTRI (3), DoRothEA (5), ChEA3-derived libraries (Literature, GTEx, ARCHS4, ENCODE, ReMap, and Enrichr) (34), five types of K562-specific regulons (S2Mb, S100Kb, S2Kb, M100Kb, and M2Kb) (9), and RegNetwork (35). Each prior was paired with established decoupleR procedures: ulm, waggr, zscore, udt, mdt, gsea, gsva, and consensus (**Figure 2B**).

**Figure 2.**
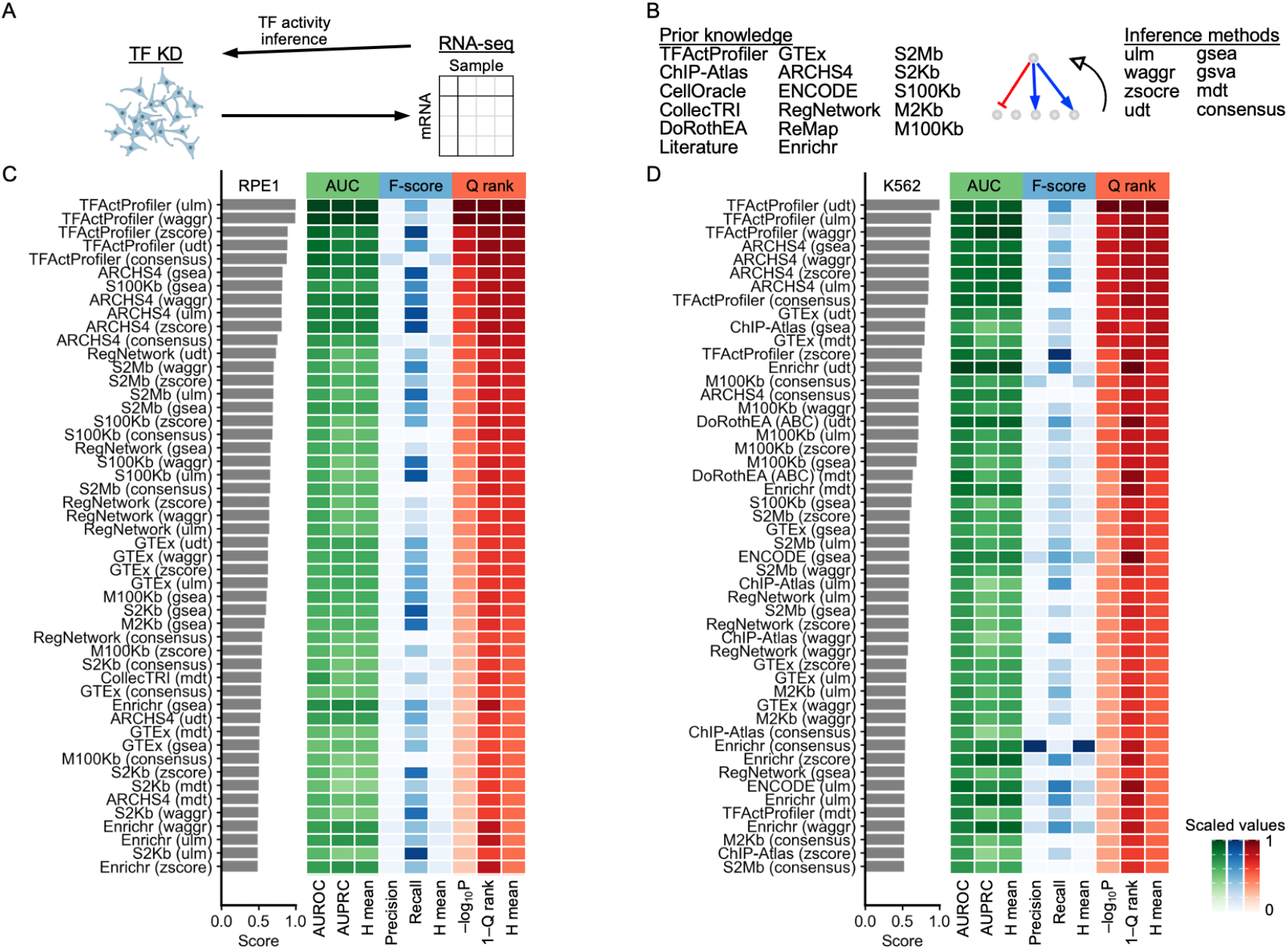
Benchmarking transcription factor (TF) activity inference across databases and methods. (**A**) Schematic of the benchmark strategy. RNA-seq profiles from individual TF knockdown (KD) experiments were used to test whether each approach correctly inferred the expected decrease in the targeted TF’s activity. (**B**) Overview of TF activity inference approaches. Combinations of prior knowledge resources (left) and inference methods (right) were compared. (**C, D**) Benchmark results in RPE1 (C) and K562 (D) cells. Bars indicate overall performance scores. Heatmaps report area under the curve (AUC), F-score, and rank-based metrics (Q rank), each scaled across methods. Only the top 50 resource–method combinations are displayed. Full results are provided in Extended Data Figure 1.

Among the tested prior-method combinations, TFActProfiler combined with ulm or udt achieved the highest overall performance across both RPE1 and K562 cells (**Figure 2C-D, Extended Data Figure 1**). Priors emphasizing breadth without functional filtering (*e*.*g*., ChIP-Atlas and CellOracle base networks) showed lower accuracy. In contrast, subset-based or curated resources (*e*.*g*., ARCHS4, GTEx, DoRothEA, and CollecTRI) outperformed the broad, unfiltered priors but did not match TFActProfiler. These results suggest that TFActProfiler provides information about the mode of regulation and effect estimates that enhance inference accuracy while maintaining broad TF and target coverage.

### Reliability estimation of TF activity inference

Inference methods that rely on prior TF–mRNA knowledge inevitably introduce error due to incomplete priors and modeling assumptions that may be violated. To quantify these deviations, we defined a mean squared error (MSE)–based reliability metric (**Figure 3A**). Higher MSE values indicate a greater violation of assumptions and reduced reliability of inferred TF activities.

**Figure 3.**
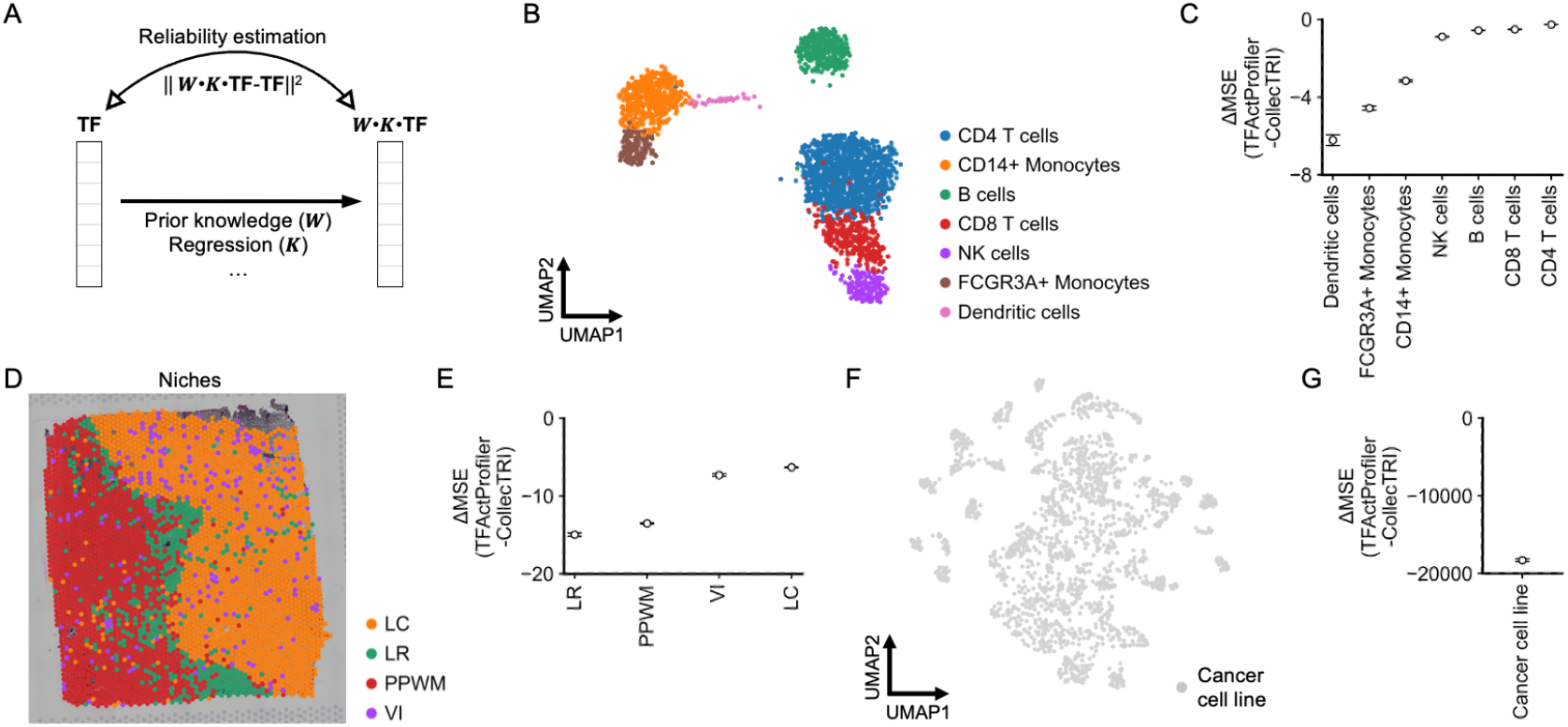
Reliability of transcription factor (TF) activity inference across cell types and spatial niches. (**A**) Framework for estimating the reliability of TF activity inference by integrating prior TF– mRNA knowledge with regression-based evaluation (Methods). (**B**) UMAP of single-cell RNA-seq profiles from peripheral blood mononuclear cells (PBMCs), annotated by cell type. (**C**) Differences in mean squared error (ΔMSE) between TFActProfiler and CollecTRI across cell types. Points indicate estimates and bars indicate 95% confidence intervals. Lower (more negative) ΔMSE indicates greater TFActProfiler reliability. (**D**) A spatial map of a chronic active multiple sclerosis lesion, segmented into a lesion core (LC), lesion rim (LR), periplaque white matter (PPWM), and perivascular infiltrate (VI). (**E**) ΔMSE between TFActProfiler and CollecTRI across niches, with 95% confidence intervals. (**F**) UMAP of bulk RNA-seq profiles from human cancer cell lines. (**G**) ΔMSE between TFActProfiler and CollecTRI across cancer cell lines.

We benchmarked TFActProfiler against CollecTRI, reportedly the most accurate resource with annotated modes of regulation (3), using single-cell RNA-seq data from peripheral blood mononuclear cells (20), spatial RNA-seq data from chronic active multiple sclerosis lesions (43), and bulk RNA-seq data from cancer cell lines (44). Across the evaluated cell types and spatial niches, TFActProfiler showed lower MSE than CollecTRI (**Figure 3B–G**). While this metric is heuristic rather than a ground-truth accuracy measure, the results align with the finding that TFActProfiler provides more reliable estimates of TF activity than CollecTRI (**Figure 2**) and support the practical utility of the MSE score for identifying unreliable TF activity estimates.

### Perturbation prediction using TF–mRNA priors

Predicting transcriptomic responses to molecular perturbations usually necessitates additional training on large-scale, single-cell, or perturbation datasets, making such approaches impractical in bulk case–control studies with limited samples. To overcome this limitation, we developed a training-free perturbation prediction framework that uses prior TF–mRNA relationships (**Figure 4A**). In both RPE1 and K562 cells, the predicted responses using TFActProfiler significantly correlated with the experimental measurements and outperformed those of CollecTRI (**Figure 4B**).

**Figure 4.**
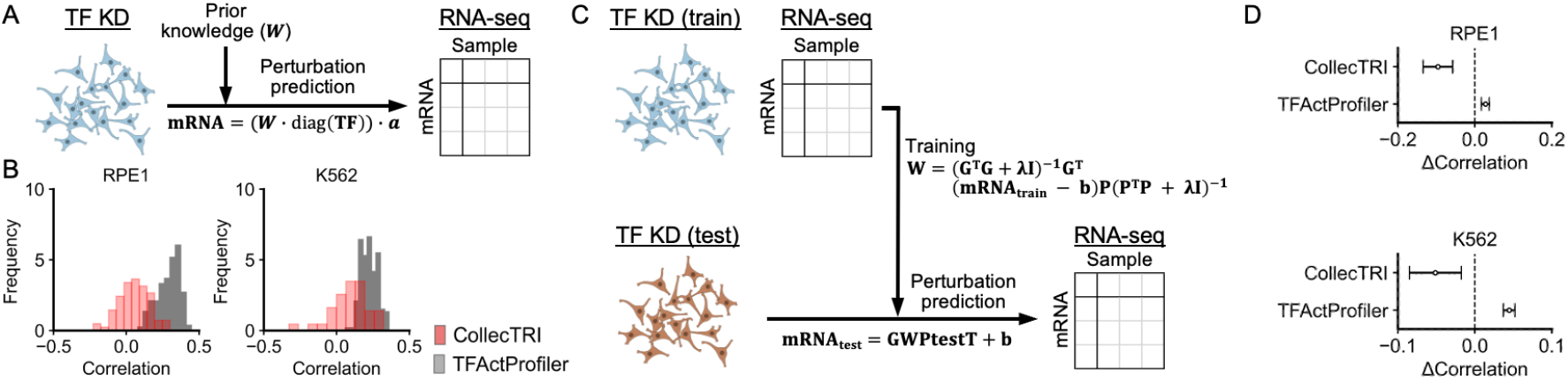
Predicting transcriptional responses to TF perturbation from TF–mRNA priors, with and without additional training. (**A**) Framework for training-free perturbation prediction. TF–mRNA prior graphs are used directly to predict gene expression changes induced by TF knockdown (KD). (**B**) Distributions of correlations between the predicted and observed mRNA expression changes in RPE1 and K562 cells. (**C**) Schematic of perturbation prediction with additional training. RNA-seq from TF KD experiments is used to fit model weights on a training set and then predict held-out perturbations. (**D**) 95% confidence intervals showing correlations between the predicted and observed expression, reported as ΔCorrelation relative to the linear model with scFoundation. Values greater than 0 indicate stronger correlations than those of the scFoundation.

Next, we examined a training-augmented setting using a linear, low-rank model. In this model, TF–mRNA priors (TFActProfiler or CollecTRI) defined the perturbation embedding and TF KD RNA-seq data provided supervision (**Figure 4C**). For comparison, we investigated scFoundation (16) embeddings with the linear model. In this held-out evaluation, TFActProfiler priors were more accurate than the linear model based on scFoundation embeddings, whereas CollecTRI was not (**Figure 4D**). Collectively, these results suggest that TFActProfiler can predict perturbation responses in a single sample without training and remains competitive with machine learning models in training-augmented frameworks.

## Discussion

This study aimed to address the longstanding trade-off between coverage and precision in TF activity inference (3, 5) and extend these estimates to *in silico* perturbation modeling. TFActProfiler integrated comprehensive TF–mRNA candidates from resources such as ChIP-Atlas and motif-based networks with transcriptomic atlases to assign modes of regulation and quantitative weights to TF–mRNA links. Since binding- or motif-only resources typically lack information about the mode of regulation, learning signed effects directly from transcriptomes reduces sign ambiguity and error propagation in regulon-based scoring. This likely explains the observed improvements in TF activity inference accuracy across perturbation benchmarks, while retaining broad coverage.

A key contribution is the uncertainty-aware readout of TF activity. Given that regulatory priors vary in quality, and assumptions such as linearity and cross-context transferability may not be valid, we defined a simple mean-squared-error-based reliability metric. This metric quantifies the self-consistency between prior-driven inference and data-driven TF–mRNA dependencies. This indicator helps to interpret activity estimates, identifies contexts where priors differ from the data, and provides a practical benchmark for comparing emerging TF–mRNA databases and inference strategies.

TFActProfiler shows that large-scale, pre-trained resources can support semi-quantitative prediction of transcriptomic responses to TF perturbations without task-specific retraining. In contrast to recent perturbation-prediction models (16, 17, 46) that require additional perturbation or single-cell data for fine-tuning, this framework remains simple, interpretable, and effective in small-sample bulk settings. When additional perturbation measurements are available, TFActProfiler’s performance was comparable to the linear model using scFoundation-derived embeddings. Of note, in benchmarks on the same human-cell datasets (6), the linear model using scFoundation-derived embeddings have been reported to outperform the mean baseline and machine learning models including scGPT (46), scBERT (47), Geneformer (48), and GEARS (49).

The current study has several limitations. Despite the improvements in coverage and accuracy, expanding to rare cell types and diverse physiological contexts requires more data and context-aware transfer strategies. Importantly, the extent to which a TF regulates a given mRNA is highly context-dependent, and our coefficients should not be interpreted as precise mechanistic truths but rather as heuristic approximations for the activity influence. Although our perturbation predictions are intended to provide directional information, they are merely linear approximations of regulatory dynamics rather than exact forecasts of expression changes. As previously described in the literature on TF activity inference and perturbation modeling (3, 17), this approach is not intended to infer a gene regulatory network (GRN). Many dedicated methods exist for GRN inference (51) and are strong alternatives for that task. Future work should integrate nonlinear or dynamic priors, refine adaptive context selection, include more detailed benchmarks (52), and extend experimental validation to determine how predicted regulatory shifts translate into phenotypic outcomes.

In conclusion, these contributions establish TFActProfiler as a resource and methodology that unites wide-coverage and uncertainty-aware TF activity inference with training-free perturbation prediction. By combining interpretability, simplicity, and general applicability, TFActProfiler provides a practical benchmark for future method development and a tool for transforming observational omics data into testable hypotheses across diverse biological settings.

## Funding

AMED-CREST (JP23gm1810009) (to EK)

Japan Society for the Promotion of Science (JSPS) KAKENHI (25K23918) (to HS)

Japan Society for the Promotion of Science (JSPS) KAKENHI (23K11312) (to KT)

## Author contributions

Conceptualization: HS, KT, ZZ, SO, TO, EK

Methodology: HS

Investigation: HS

Visualization: HS

Project administration: EK

Supervision: EK

Writing – original draft: HS

Writing – review & editing: KT, ZZ, SO, TO, EK

## Conflict of interest

Authors declare that they have no competing interests.

## Data availability

The RNA-seq datasets from RPE1 and K562 cell lines are available from a previous study (PRJNA831566) (31). The peripheral blood mononuclear cell (PBMC) dataset and the chronic active multiple sclerosis lesion dataset were distributed with decoupleR (v2.1.1). The expression dataset of human cancer cell lines was downloaded from the DepMap portal website (https://depmap.org/portal). The TFActProfiler code is available on our GitHub repository (https://github.com/HikaruSugimoto/tfactprofiler).

---

**Extended Data Figure 1.**
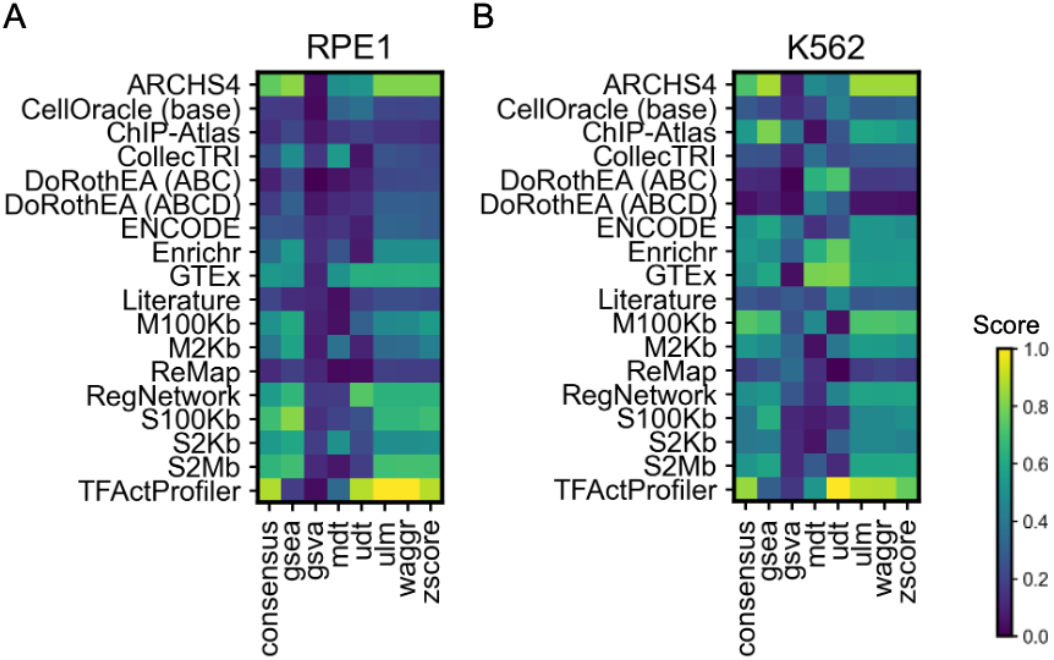
Benchmarking transcription factor (TF) activity inference across databases and methods. Heatmaps summarizing overall performance scores for RPE1 (**A**) and K562 (**B**) cells. Rows correspond to TF–target databases and columns to inference methods. Colors indicate the aggregate performance score (0–1; higher is better), using the same scale in both panels.

## Notes

### Competing Interest Statement

The authors have declared no competing interest.

